# Dynamic control over feedback regulatory mechanisms improves NADPH fluxes and xylitol biosynthesis in engineered *E. coli*

**DOI:** 10.1101/2020.07.27.222588

**Authors:** Shuai Li, Eirik A. Moreb, Zhixia Ye, Jennifer N. Hennigan, Daniel Baez Castellanos, Tian Yang, Michael D. Lynch

## Abstract

We report improved NADPH flux and xylitol biosynthesis in engineered *E. coli*. Xylitol is produced from xylose via an NADPH dependent reductase. We utilize two-stage dynamic metabolic control to compare two approaches to optimize xylitol biosynthesis, a stoichiometric approach, wherein competitive fluxes are decreased, and a regulatory approach wherein the levels of key regulatory metabolites are reduced. The stoichiometric and regulatory approaches lead to a 16 fold and 100 fold improvement in xylitol production, respectively. Strains with reduced levels of enoyl-ACP reductase and glucose-6-phosphate dehydrogenase, led to altered metabolite pools resulting in the activation of the membrane bound transhydrogenase and a new NADPH generation pathway, namely pyruvate ferredoxin oxidoreductase coupled with NADPH dependent ferredoxin reductase, leading to increased NADPH fluxes, despite a reduction in NADPH pools. These strains produced titers of 200 g/L of xylitol from xylose at 86% of theoretical yield in instrumented bioreactors. We expect dynamic control over enoyl-ACP reductase and glucose-6-phosphate dehydrogenase to broadly enable improved NADPH dependent bioconversions.

**Highlights:**

- Decreases in NADPH pools lead to increased NADPH fluxes
- Pyruvate ferredoxin oxidoreductase coupled with NADPH-ferredoxin reductase improves NADPH production *in vivo*.
- Dynamic reduction in acyl-ACP/CoA pools alleviate inhibition of membrane bound transhydrogenase and improve NADPH flux
- Xylitol titers > 200g/L in fed batch fermentations with xylose as a sole feedstock.

## Introduction

Xylitol in an industrial sugar alcohol with a primary use as a sweetener, ^1^ and is produced at ~125,000 tons annually, via the oxidation of xylose. ^1–3^ Xylose is the second most abundant natural sugar (after glucose), which can be isolated from hemicellulose. ^4^ Many studies have demonstrated the use of xylose as a feedstock for the biosynthesis of numerous products ranging from biofuels (ethanol) to numerous chemicals including lactic acid, succinic acid, xylonate, 1,2,4-butanetriol, and xylitol. ^5–8^ Perhaps the simplest conversion is xylose to xylitol, which requires only a single enzyme, a xylose reductase and a reductant (NADPH). Biosynthetic production of xylitol has the potential to decrease costs, while avoiding the use of organic solvents and eliminating the need for expensive reduction catalysts. ^9^ Most previous studies producing xylitol from xylose rely on a bioconversion requiring an additional sugar (usually glucose) as an electron donor. ^3,10,11^ Oxidation of glucose (producing the byproduct gluconic acid) generates NADPH which is then used for xylose reduction.^12^ While these processes offer high xylitol titers and a good yield when considering xylose, the requirement for glucose at equimolar levels to xylose is a significant inefficiency. More broadly, improving NADPH availability or flux, useful in the synthesis of numerous metabolites as well as cell based bioconversions, has been a long standing challenge in metabolic engineering. ^13–17^

We sought to apply two-stage dynamic metabolic control (DMC) to improve NADPH flux and xylitol production using xylose as a sole feedstock.^18^ Dynamic control over metabolism has become a popular approach in metabolic engineering, and has been used for the production of various products from 3-hydroxypropionic acid to myo-inositol and many others. ^19–22^ We have recently reported an extension of dynamic metabolic control to two-stage bioprocesses, where products are made in a metabolically productive phosphate depleted stationary phase. ^23,24^ The implementation of this approach relies on combined use of controlled proteolysis and gene silencing, using degron tags and CRISPR interference respectively. ^25–27^ Importantly, in these initial studies we demonstrated that improved metabolic fluxes resulting from dynamic metabolic control, can be a consequence of reducing levels of central metabolites which are feedback regulators of other key metabolic pathways.^24^ Specifically, we have recently shown that decreasing glucose-6-phosphate dehydrogenase levels activates the SoxRS regulon increasing expression and activity of pyruvate ferredoxin/flavodoxin oxidoreductase (Pfo). Pfo leads to improved acetyl-CoA production in stationary phase. (Refer to Figure 3a). ^24^ In this work we report the evaluation of combinations of synthetic metabolic valves on xylitol production from xylose. Firstly, increased Pfo activity not only leads to improved acetyl-CoA flux but also NADPH production. NADPH is produced from reduced flavodoxin/ferredoxin via the action of NADPH dependent flavodoxin/ferredoxin reductase (Fpr). We also identify a key regulatory mechanism controlling NADPH fluxes, namely the inhibition of the membrane bound transhydrogenase (PntAB) by fatty acid metabolites. By dynamically disrupting fatty acid biosynthesis, we alleviate inhibition of PntAB. This mechanism is synergistic with activating Pfo and greatly increases NADPH flux and xylitol production. We compare this “regulatory” approach with a more intuitive stoichiometric strategy where the levels of key enzymes competing with xylitol production are dynamically reduced. Importantly, improved NADPH fluxes are, in part, a consequence of reduced NADPH pools. Reduced NADPH pools drive changes in expression and activity that result in increased NADPH fluxes, presumably a regulatory mechanism which has evolved to restore set point NADPH levels. These results are a reminder that pools and flux are not equivalent and not necessarily correlated.

## Results

### Stoichiometric Strategy

We initially rationally designed strains to optimize xylitol production from xylose utilizing two stage dynamic metabolic control, reliant on decreasing levels of key competitive pathways. As illustrated in Figure 1a, this design included dynamic reduction in xylose isomerase (*xylA*) and soluble transhydrogenase (*udhA*) activities. These modifications were designed to reduce xylose metabolism which competes with xylitol production and increases NADPH supply. NADPH can be consumed by the soluble transhydrogenase. ^28^ Toward this goal we constructed strains and plasmids to enable the dynamic reduction in XylA and UdhA levels upon phosphate depletion.^29,30^ Refer to Supplemental Table S1 for strains and plasmids used in this study. As described above and previously reported dynamic reduction in activity was accomplished by adding C-terminal DAS+4 degron tags to the chromosomal *xylA* and *udhA* genes as well as the overexpression of guide RNAs aimed at silencing their expression.^24–26^ The impact of these modifications on enzyme levels in two stage cultures is given in Figures 1b-c. In the case of XylA, proteolysis led to ~ 60% reduction in activity. To our surprise the silencing gRNA actually led to an increased XylA activity level. The mechanism behind this is currently unknown and requires further study. The combination of silencing and proteolysis resulted in no further reduction in activity when compared to proteolysis alone. In the case of UdhA, proteolysis resulted in a ~30% reduction in activity, whereas silencing had no detectable impact on activity levels with or without proteolysis.

**Figure 1:**
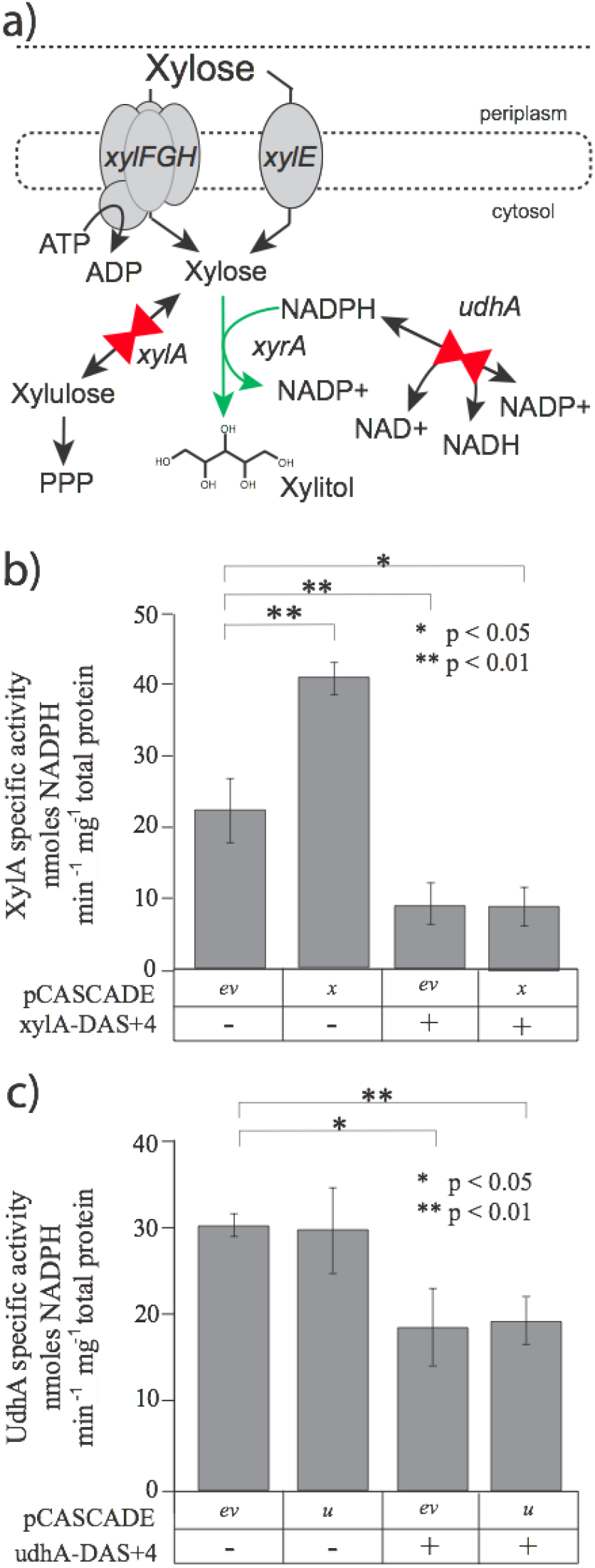
a) A stoichiometric approach to improving xylitol production using dynamic control, wherein enzyme levels of competitive pathways are dynamically reduced to redirect flux to the desired product. In this case xylose isomerase (XylA) and the soluble transhydrogenase (UdhA) were targeted for dynamic control. Enzyme levels of b) XylA and c) UdhA in response to induble proteolysis and/or gene silencing in a phosphate depleted stationary phase. ev -empty vector, x- *xylA* promoter, u- *udhA* promoter.

The combination of proteolysis and silencing for XylA or “X valves” and proteolysis alone in the case of UdhA, a “U Valve”, were evaluated for xylitol production. Specifically strains were engineered with these metabolic valves as well as for overexpression of a xylose reductase (*xyrA* from *A. niger*).^31,32^ and evaluated in two-stage minimal media microfermentations as reported by Moreb et al. ^29^ Results are given in Figure 2a-c. Additionally, a confirmatory analysis of XyrA kinetics was performed, and results are given in Supplemental Figure S1. The combination of modifications resulted in a 16 fold increase in xylitol production compared to the control.

**Figure 2:**
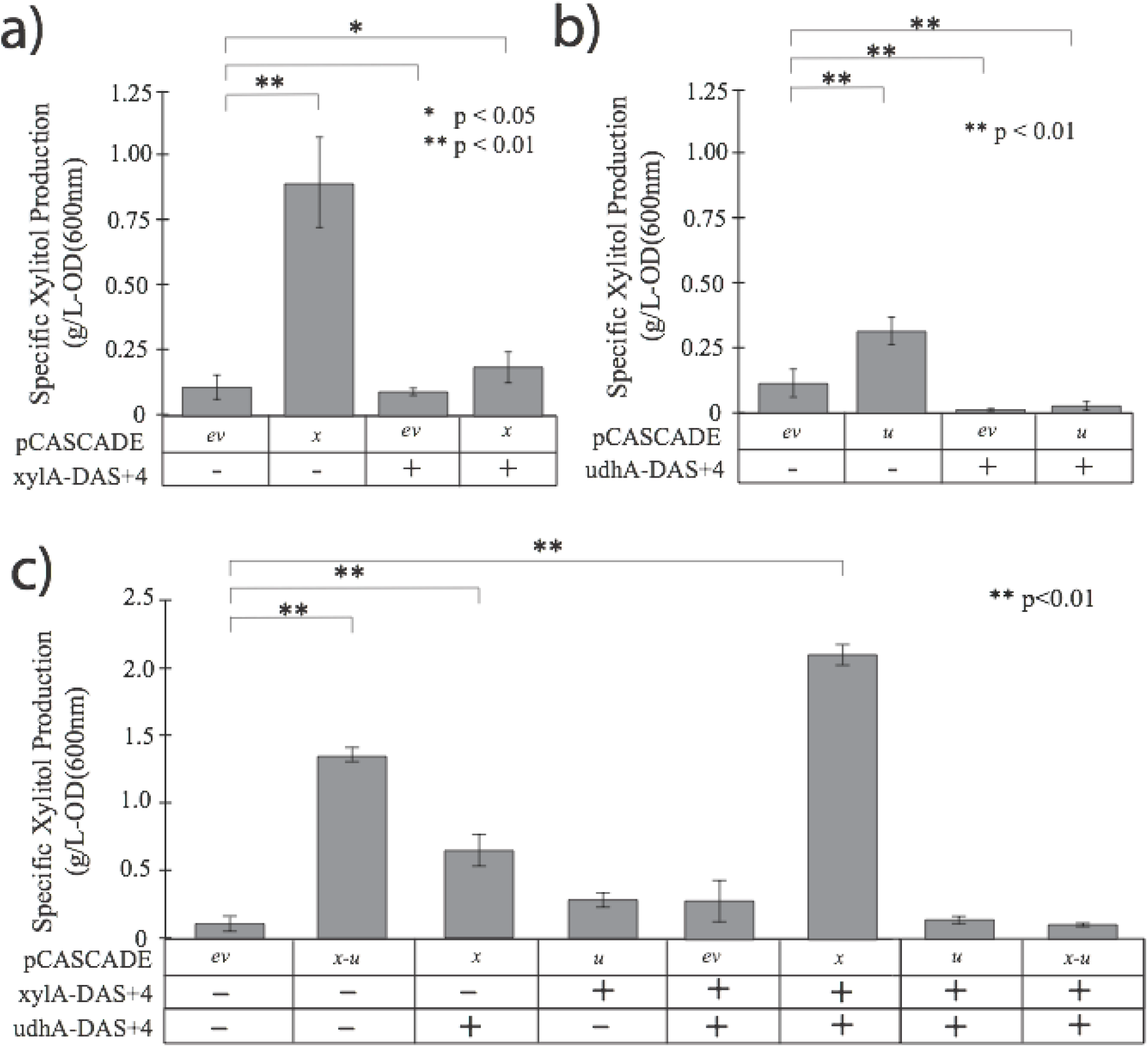
Specific xylitol production in strains engineered for dynamic control over levels of a) xylose isomerase (XylA), b) soluble transhydrogenase (UdhA) and c) the combined control over xylose isomerase soluble transhydrogenase. ev -empty vector, x- *xylA* promoter, u- *udhA* promoter. All results were obtained from microfermentations.

### Regulatory Strategy

To investigate the impact of a regulatory strategy, we next sought to evaluate the potential impact of a larger set of valves on xylitol production as illustrated in Figure 3a. We constructed a set of strains with valves in citrate synthase (GltA), glucose-6-phosphate dehydrogenase (Zwf) and enoyl-ACP reductase (FabI) which control flux through the tricarboxylic acid cycle, pentose phosphate pathway and fatty acid biosynthesis, respectively. We have previously reported the construction of metabolic valves in GltA (“G Valves”), and Zwf (“Z Valves”) which comprised either proteolytic degradation (DAS+4 tags), gene silencing (either the zwf promoter or gltAp2 promoter) or both. ^23,25,26^ In the case of FabI, we constructed new strains and plasmids to evaluate two-stage dynamic control on FabI levels. Toward this goal, as similarly reported by Li et al., ^24^ we appended a superfolder GFP to the C-terminus of the *fabI* allele to enable quantification of protein levels by an ELISA. Unfortunately and unexpectedly, when plasmids silencing *fabI* expression were evaluated, guide RNA protospacer loss was observed (Supplemental Figure S2) and as a result we could not reliably obtain results where *fabI* is silenced. FabI proteolysis led to a ~75 % reduction in FabI levels (Supplemental Figure S3), and as a result proteolytic degradation alone will be referred to as an “F Valve”. Strains were constructed with combinations of “X”, “U”, “G”, “Z” and “F” valves and evaluated for xylitol production, again in minimal media microfermentations. Results are given in Figure 3. To our surprise the highest xylitol producer had neither “X” or “U” valves but rather a combination of “F” and “Z” valves. Xylitol production in the “FZ” valve strain was synergistic above either “F” or “Z” valves alone (Figure 3b). This was surprising in that these two enzymes have no direct or predictable impact of xylitol biosynthesis as can be seen in Figure 3a. We have recently reported that the “Z” valve results in increased acetyl-CoA fluxes by leading to the activation of Pfo (encoded by the *ydbK* gene).^24,33^ With increased flux through Pfo we hypothesized that NADPH could be generated from reduced ferredoxin/flavodoxin through ferredoxin reductase (Fpr). Using deletions in *ydbK* and *fpr*, as shown in Figure 4a, we confirmed that this pathway, and specifically Fpr, is indeed in part responsible for the elevated xylitol production and NADPH flux observed in our “FZ” valve strain. This is, to our knowledge, the first confirmation that Fpr is reversible *in vivo*. This reverse flux through Fpr may be dependent on low NADPH pools (as discussed below). The synergistic impact of the “F” valves was somewhat unanticipated. However, elevated NADPH fluxes due to dynamic control over FabI (enoyl-ACP reductase) can be attributed to reduced levels of fatty acid metabolites, specifically acyl-CoAs (and potentially their precursors acyl-ACPs). Fatty acyl-CoAs are competitive inhibitors of the membrane bound transhydrogenase encoded by the *pntAB* genes (Figure 3a).^34–36^ Palmitoyl-CoA, specifically, has a reported Ki of 1-5 *μ*M.^34–36^ Control over FabI levels and/or activity has been previously shown to reduce acyl-ACP pools and as a result alleviate feedback inhibition of acetyl-CoA carboxylase and malonyl-CoA synthesis.^37–41^ To our knowledge this is the first study demonstrating the importance of these metabolites in controlling NADPH fluxes. While previous reports demonstrate the inhibition of PntAB by acyl-CoAs (which in minimal media are derived from fatty acid biosynthesis),^34–36^ there remains a possibility that acyl-ACPs also act as inhibitors, although future work is needed to confirm this hypothesis.

**Figure 3:**
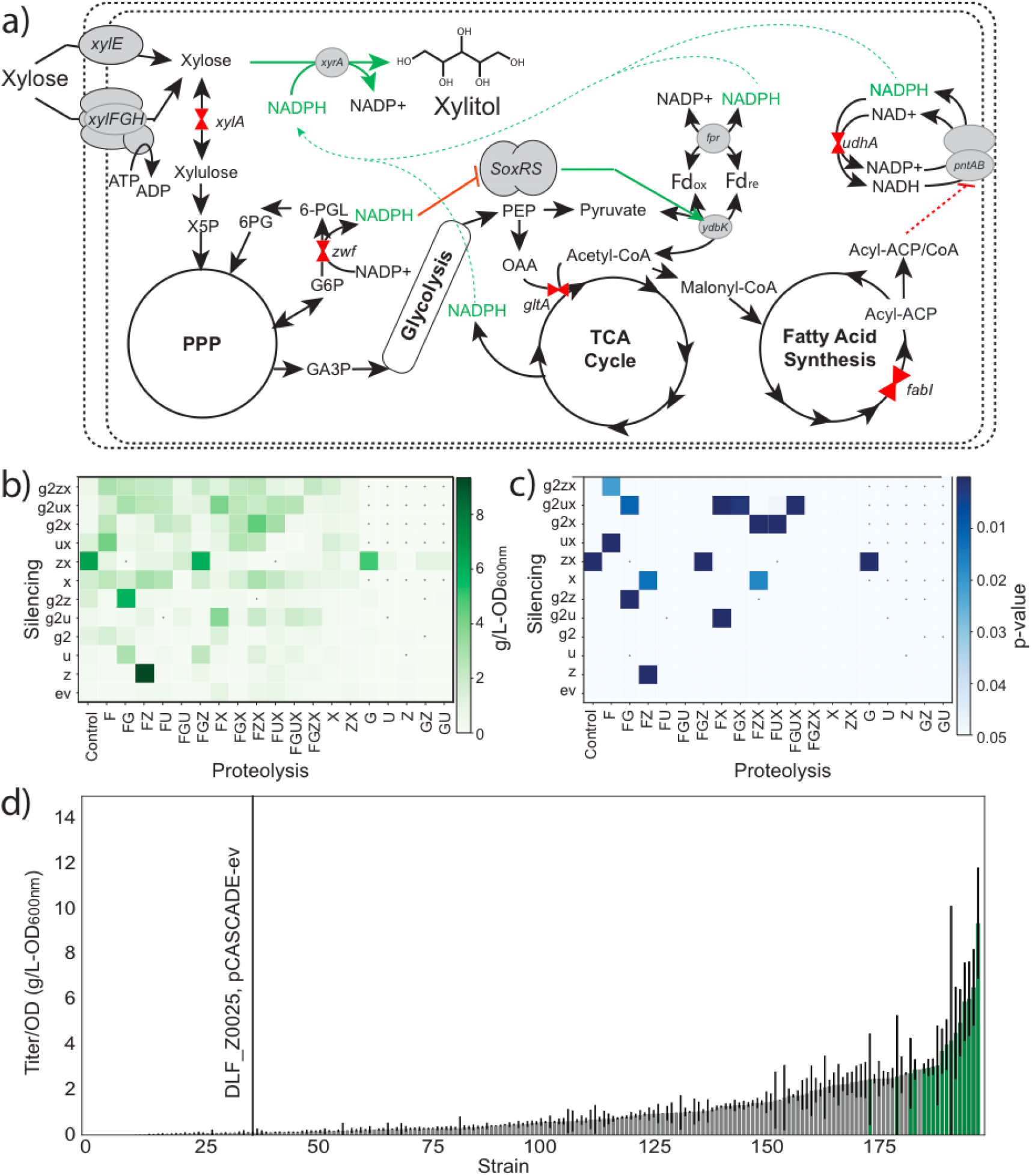
a) An overview of xylitol production and the location of metabolic valves in central metabolism. Xylitol is produced from xylose by a xylose reductase (*xyrA*). Valves comprise inducible proteolysis and/or silencing of 5 enzymes: citrate synthase (gltA), xylose isomerase (xylA), glucose-6-phosphate dehydrogenase (zwf), enoyl-ACP reductase (fabI) and soluble transhydrogenase (udhA). The membrane bound transhydrogenase (pntAB) is also shown. b) Specific xylitol production (g/L-OD_600nm_) in microfermentations as a function of silencing and or proteolysis. c) P-values for the data in b, comparing each strain to the no-valve control using a Welchs t-test. d) a rank order plot of the data from panel. Green bars indicate a p-value < 0.05. Abbreviations: *xylE* : xylose permease, *xylFGH* : xylose ABC transporter, PPP: pentose phosphate pathway, PDH: pyruvate dehydrogenase multienzyme complex, TCA: tricarboxylic acid, G6P: glucose-6-phosphate, 6-PGL: 6-phosphogluconolactone, 6PG: 6-phosphogluconate, GA3P: glyceraldehyde-3-phosphate, PEP: phosphoenolpyrvate, OAA: oxaloacetic acid, X5P: xylulose-5-phosphate, Fd: ferredoxin. Silencing: ev: empty vector, g2: *gltAp2* promoter, z: *zwf* promoter, x: *xylA* promoter, u: *udhA* promoter. Proteolysis: F: fabI-DAS+4, G: gltA-DAS+4, Z: zwf-DAS+4, U: udha_DAS+4, X: xylA-DAS+4. All results were obtained from microfermentations.

**Figure 4 :**
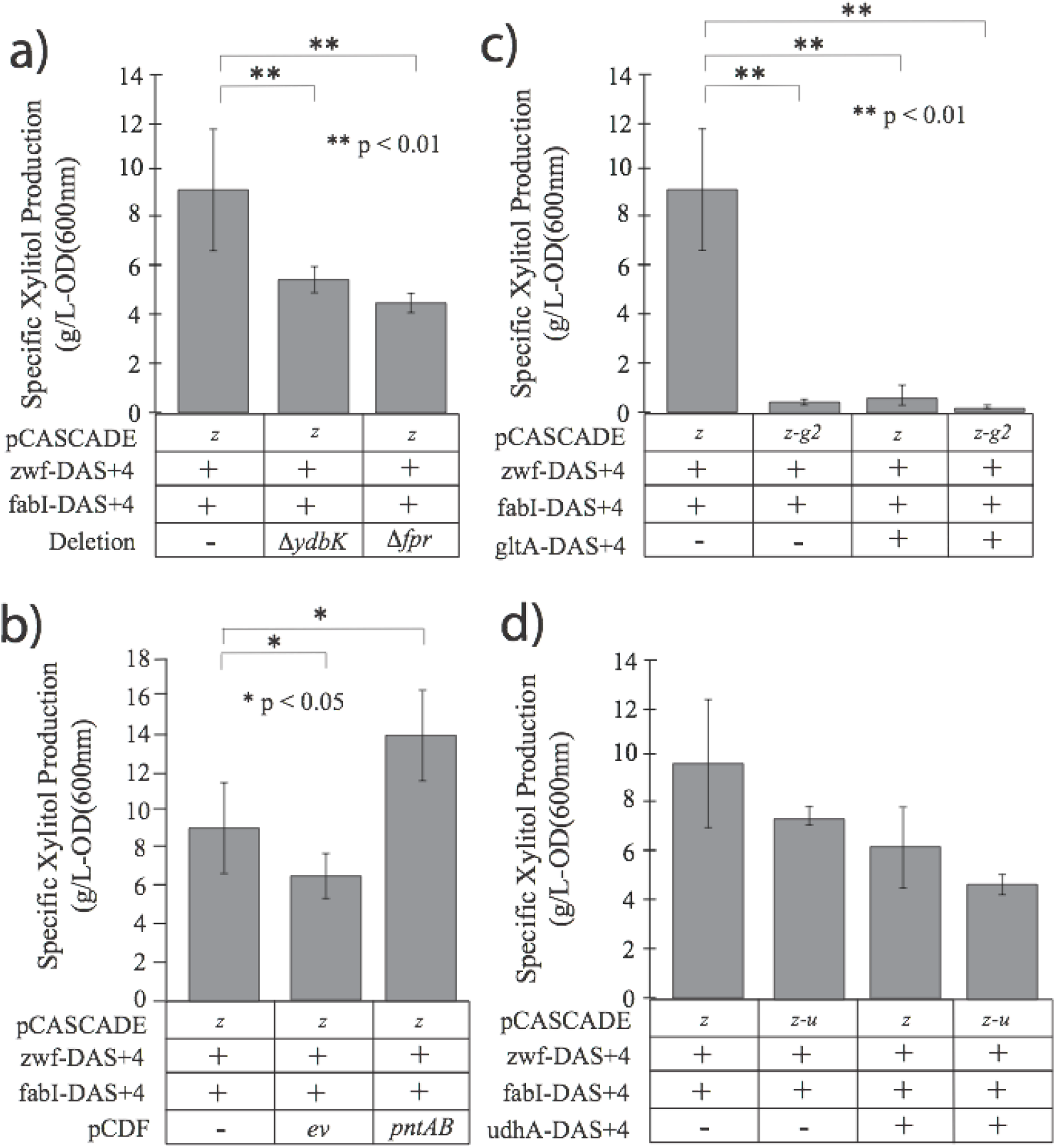
Identification of pathways responsible for NADPH and xylitol production in the “FZ” valve strain a) the impact of deletions of ydbK and fpr on specific xylitol production, b) the impact of pntAB overexpression on xylitol production. (c-d) “FZ” valve strains further modified for dynamic control over c) GltA levels and d) UdhA levels. ev -empty vector, z- *zwf* promoter, g2- *gltAp2* promoter, u- *udhA* promoter. All results were obtained from microfermentations.

We next evaluated several additional modifications on top of the “FZ” valves, with a potential to impact xylitol production. (Figure 4b-d). Specifically we evaluated the addition of “G” and “U” valves as well as overexpression of pntAB. Plasmid based overexpression of the *pntAB* genes (using a low phosphate inducible promoter^29^) led to a significant improvement in xylitol production (Figure 4b). In contrast, the addition of either the “G” or “U” valve to the “FZ” combination did not increase xylitol synthesis but rather led to a significant decrease in xylitol production (Figure 4c-d). This suggests that citrate synthase (GltA) activity, and flux through the TCA cycle, is required for optimal NADPH flux.

Using results from these experiments, we were able to estimate boundary conditions for several intracellular fluxes. For example from Figure 4b, we can estimate that flux through the Pfo/Fpr pathway accounts for at most ~ 55% of the NADPH/xylitol production. As a result we are able to build stoichiometric metabolic models, as illustrated in Figure 5, comparing an optimal growth phase and xylitol production phase.^42,43^ Importantly, this model confirms that indeed TCA flux is critical for xylitol production (Supplemental Figure S4) and that a 4-fold increase in PntAB activity, in addition to flux through the Pfo/Fpr pathways is needed to explain increases in NADPH flux and xylitol production. The model predicts an overall maximal xylitol yield in this metabolic state of ~0.864g/g of xylose, in line with yields measured in fed batch fermentations as discussed below.

**Figure 5:**
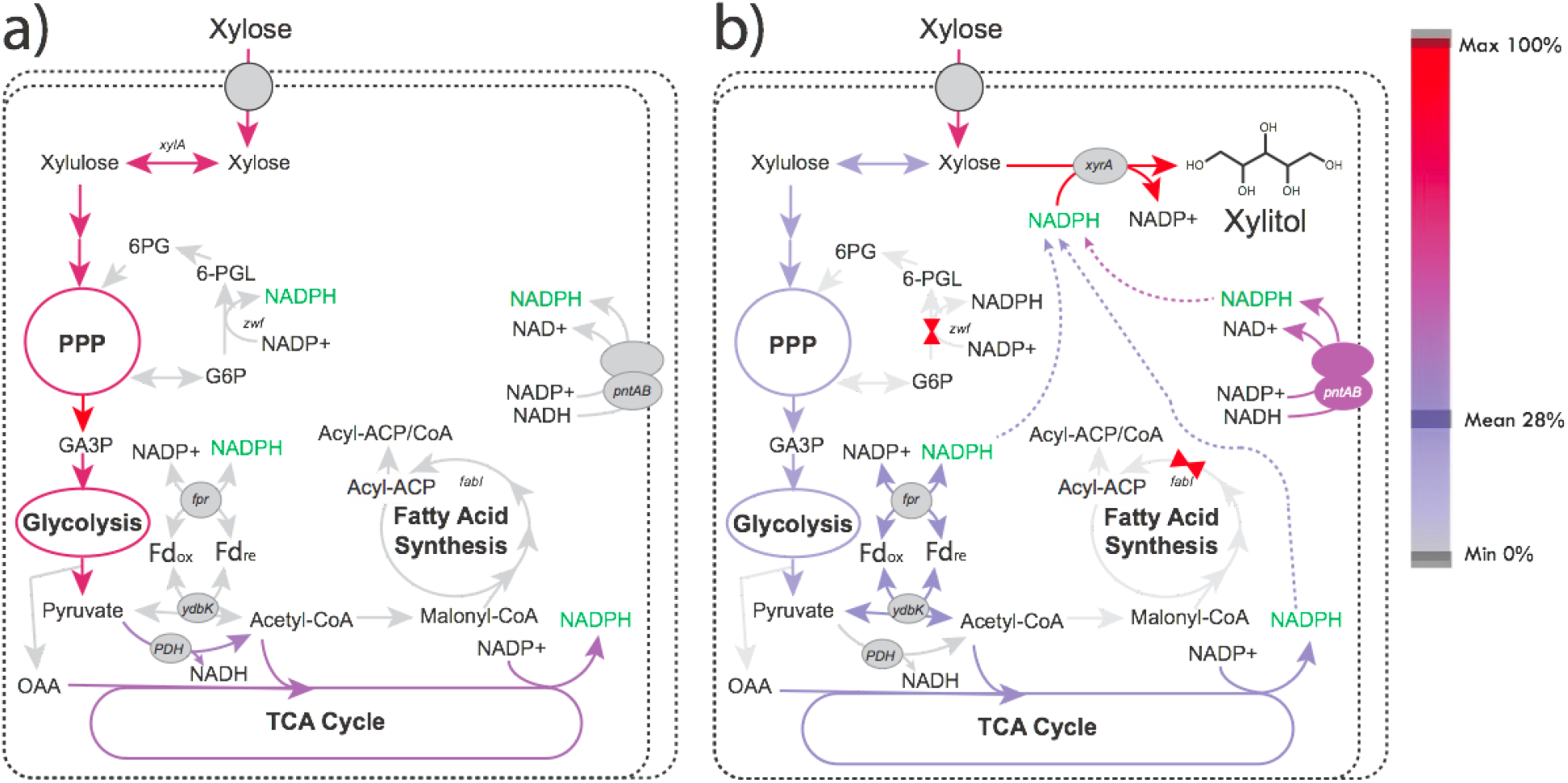
Stoichiometric flux models of a) cellular growth and b) stationary phase xylitol production in “FZ” valve strains. Pathway flux is relative to xylose uptake rates. During growth the majority of flux is through the pentose phosphate pathway (PPP), pyruvate dehydrogenase multienzyme complex (PDH) with minimal flux through the pentose membrane bound transhydrogenase. Upon dynamic control, a 4-fold increase in membrane bound transhydrogenase flux is accompanied by increased flux through Pfo (ydbK) and FPr. Abbreviations: G6P: glucose-6-phosphate, 6-PGL: 6-phosphogluconolactone, 6PG: 6-phosphogluconate, GA3P: glyceraldehyde-3-phosphate, OAA: oxaloacetic acid.

### Production in Instrumented Bioreactors

Next, we compared xylitol production in instrumented bioreactors using the “FZ” valve strain with and without pntAB overexpression with a control strain. Minimal media fed batch fermentations were performed as described by Menacho-Melgar et al.,^30^ wherein the media has enough batch phosphate to support target biomass levels (~ 25gCDW/L) prior to phosphate depletion and induction of xylitol biosynthesis in stationary phase. Results of these studies are given in Figure 6. While *xyrA* expression in our control strain (DLF_Z0025) led to only a few grams per liter of xylitol (Figure 6a), the incorporation of “FZ” valves led to titers over 100g/L in 160 hours of production (Figure 6b). The additional overexpression of *pntAB* (Figure 6c) resulted in maximal titers over 200g/L (185-204g/L) in 170 hrs. In these duplicate fermentations the average overall xylitol yield was 0.873 +/− 0.026 g/g xylose, and the average production yield (in stationary phase) was 0.935 +/− 0.011 g/g xylose.

**Figure 6:**
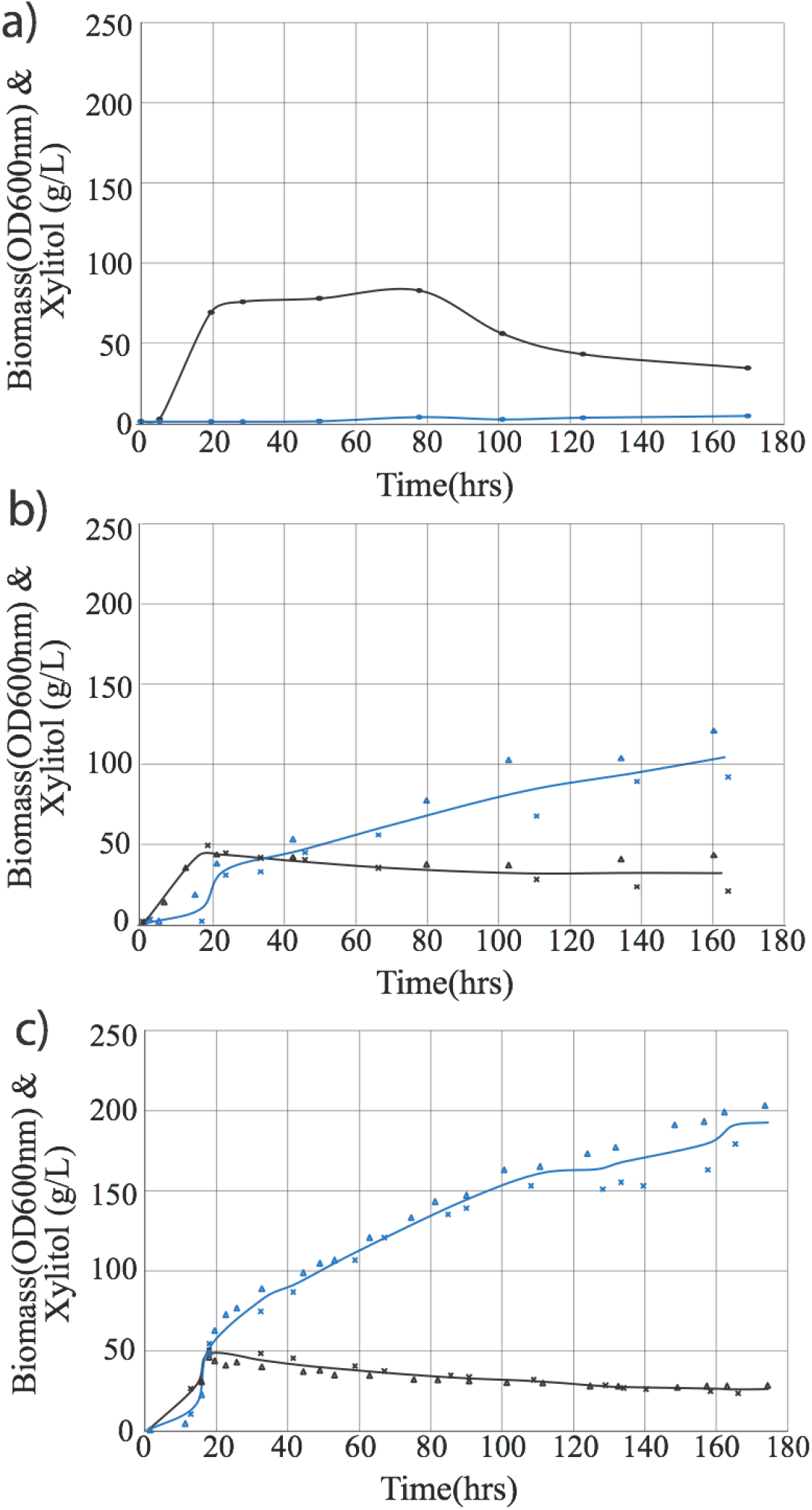
Xylitol production in minimal media fed batch fermentations in instrumented bioreactors by a) the control strain expressing xylose reductase (DLF_Z0025, pCASCADE-ev, pHCKan-xyrA), b) the “FZ” valve strain (DLF_Z0025-fabI-DAS+4-zwf-DAS+4, pCASCADE-z, pHCKan-xyrA) c) the “FZ” valve strain also overexpressing the membrane bound transhydrogenase *pntAB* (DLF_Z0025-fabI-DAS+4-zwf-DAS+4, pCASCADE-z, pHCKan-xyrA, pCDF-pntAB). Biomass (black) and xylitol (blue) are given as a function of time. For b) & c) x’s and triangles represent the measured values of two duplicate runs.

### Improved NADPH Flux is not Correlated with NADPH pools

Lastly, we measured the levels of NADPH in a set of our engineered strains. Results are given in Figure 7a. Interestingly, there was no correlation between specific xylitol production and NADPH pools. In this case, the three strains having the highest NADPH pools were the control strain and the strains with dynamic control over enoyl-ACP levels (“F” valve) or soluble transhydrogenase (“U” valve) levels. The addition of the “Z” valve (reduced levels of glucose-6-phosphate dehydrogenase) led to a decrease in NADPH pools but an increase in NADPH flux. Deletions of either *ydbK* and or *fpr*, also led to decreases in NADPH levels, and while overexpression of *pntAB* increased xylitol production rates and fluxes it did not improve NADPH pools in the “FZ” background.

**Figure 7:**
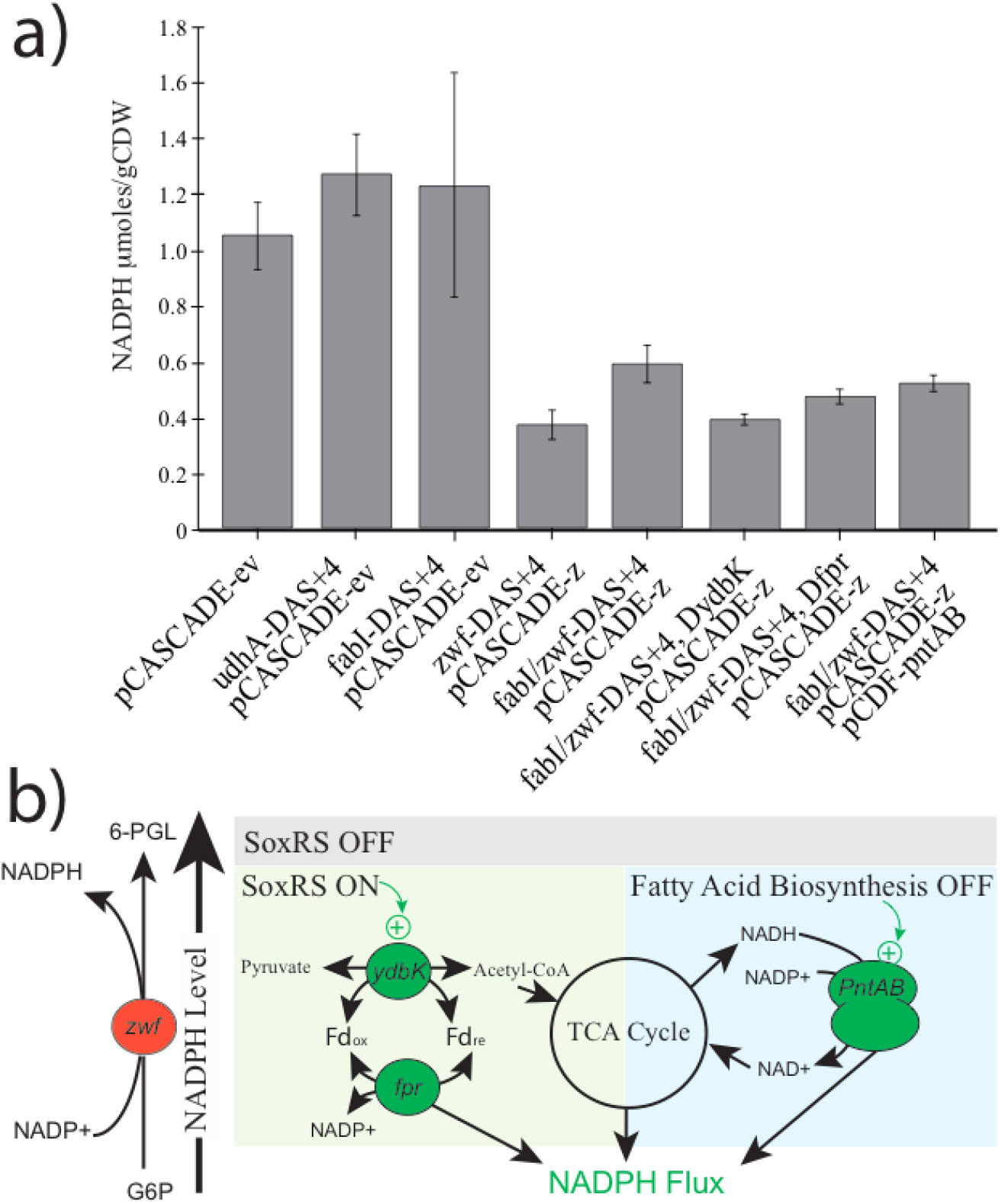
a) Stationary phase NADPH pools in engineered strain. Pools were measured 24 hours post phosphate depletion. b) A conceptual model of two-stage NADPH production in our engineered system. Glucose-6-phosphate dehydrogenase (encoded by the *zwf* gene) is normally responsible for the biosynthesis of a majority of NADPH. This irreversible reaction drives an NADPH set point, in which the SoxRS oxidative stress response is OFF (gray area). Dynamic reduction in Zwf levels reduces NADPH pools activating the SoxRS response, which in turn activates expression of Pyruvate ferredoxin oxidoreductase (Pfo, encoded by the *ydbK* gene) and NADPH dependent ferredoxin reductase (Fpr). Together Pfo and Fpr (operating in reverse) constitute a new pathway to generate NADPH as well as allow for continued pyruvate oxidation and generation of acetyl-CoA for entry into the tricarboxylic acid cycle (TCA cycle). NADPH flux is further enhanced by reducing fatty acid biosynthesis whose products inhibit the membrane bound transhydrogenase (encoded by the pntAB genes). Activated PntAB uses the proton motive force to convert NADH from the TCA cycle to NADPH. NADPH can be used for bioconversions such as for xylitol production.

## Discussion

The use of 2-stage dynamic control generated an usual metabolic state leading to enhanced NADPH fluxes and xylitol production. To our knowledge this is the highest titer and yield of xylitol produced to date in engineered *E. coli,* particularly with xylose as a sole carbon source. Additionally, the productive stationary phase generated with these modifications can be extended to at least 170 hours. While the focus of this work has been on xylitol production, the identification of “F” and “Z” valves impacting NADPH flux has applicability to other NADPH dependent processes including more complicated pathways, and may represent a facile method for routine NADPH dependent bioconversions. ^44–46^ The impact of FabI activity and fatty acid metabolite pools, on transhydrogenase activity, is consistent with previous biochemical studies, ^34–36^ and has likely evolved to balance NADPH supply with fatty acid synthesis demand. Unfortunately, this feedback regulatory mechanism has been lost in the past several decades of metabolic engineering studies in *E. coli*, ^47^ yet represents a powerful approach to improving NADPH fluxes. The unpredictable combination of “F” and “Z” valves is at odds with standard thinking regarding NADPH flux, where Zwf is often considered one of the primary sources of NADPH in the cell and reducing Zwf activity would not be high on a list of changes to make in order to increase NADPH supply.^48,49^

In order to explain the lack of correlation between NADPH pools and our results, we developed a conceptual model as illustrated in Figure 7b. The “Z” valve leads to a decrease in NADPH pools which activate the SoxRS regulon, which is sensitive to oxidant and NADPH levels. ^33,50^ SoxRS activation leads to increased expression of Pfo, which is required to maintain a high rate of pyruvate oxidation, generating NADPH via Fpr.^33,51,52^ Uniquely, this study identifies a previously unreported pathway for NADPH production utilizing Pfo and Fpr and supports that Fpr catalyzes a reversible reaction *in vivo*. Pfo expression is required, not only for pyruvate oxidation and sugar consumption but also NADH generation via the TCA cycle. Increased TCA flux produces excess NADH which is needed as a substrate for PntAB for maximal NADPH flux. Disruption of the TCA cycle (“G” Valve, Figure 4c) eliminates NADH production and acetyl-CoA consumption, greatly reducing NADPH flux. Increased NADPH levels due to the “F” valve make sense in light of the results discussed and are attributable to increased activity of the membrane bound transhydrogenase, PntAB. Reduced soluble transhydrogenase (UdhA, Figure 4d) levels leads to increased NADPH pools (Figure 7a) which presumably reduce SoxRS activation and Pfo expression. Simply put, the metabolic network responds to decreased NADPH and acyl-CoA pools by increasing sugar consumption and NADPH flux to compensate. If “set” point NADPH pools are regained or if continued sugar catabolism stops, continued NADPH flux is halted.

Lastly, the metabolic state leading to enhanced NADPH flux and xylitol production would be hard to identify and/or engineer in a growth coupled process as it relies on the manipulation of feedback inhibition due to central metabolites. These central metabolic regulatory circuits have evolved to balance fluxes to both optimize growth and enable adaptive responses to environmental and physiological perturbations. Dynamic metabolic control, and in particular two-stage dynamic metabolic control, is uniquely suited to manipulate central metabolite levels without impacting cell growth or survival. This approach can lead to the discovery as well as the manipulation of central regulatory mechanisms, which in turn have a high potential to enhance metabolic fluxes and drive future metabolic engineering strategies.

## Materials & Methods

### Reagents and Media

All reagents and chemicals were obtained from Sigma Aldrich (St. Louis, MO) unless otherwise noted. MOPS (3-(N-morpholino)propanesulfonic acid) was obtained from BioBasic, Inc. (Amherst, NY). Crystalline xylose was obtained from Profood International (Naperville, IL). All media: SM10++, SM10 No Phosphate, and FGM25 were prepared as previously reported,^30^ except that xylose was substituted for glucose (1 gram xylose for 1 gram glucose) in all media formulations. LB, Lennox formulation, was used for routine strain propagation. Working antibiotic concentrations were as follows: kanamycin: 35 μg/mL, chloramphenicol: 35 μg/mL, gentamicin:10 μg/mL, 10 zeocin: 100 μg/mL, blasticidin: 100 μg/mL, spectinomycin: 25 μg/mL, tetracycline: 5 μg/mL.

### Strains & Plasmids

Refer to Supplemental Table S1 for a list of strains and plasmids used in this study. Sequences of synthetic DNA used in this study are given in Supplemental Table S2. Chromosomal modifications were constructed using standard recombineering methodologies. ^53^ The recombineering plasmid pSIM5 was a kind gift from Donald Court (NCI, https://redrecombineering.ncifcrf.gov/court-lab.html). ^53,54^ C-terminal DAS+4 tags were added by direct integration and selected through integration of antibiotic resistance cassettes 3’ of the gene as previously described.^24^ All strains were confirmed by PCR, agarose gel electrophoresis and confirmed by sequencing. Refer to Supplemental Table S3 for oligos used for strain confirmation and sequencing.

The *xyrA* gene from *Aspergillus niger* was codon optimized for expression in *E. coli* and the plasmid, pHCKan-xyrA (Addgene #58613), enabling the low phosphate induction of xylose reductase, was constructed by TWIST Biosciences (San Francisco, CA). pCDF-pntAB (Addgene # 158609) was constructed using PCR and Gibson Assembly from pCDF-ev ^30^ to drive expression of the *pntAB* operon from the low phosphate inducible ugpBp promoter. ^29^. pCASCADE guide RNA array plasmids were prepared by the combination of PCR and Gibson assembly as previously described.^24^ Refer to Supplemental Table S4 for oligos used for pCASCADE plasmid construction.

### Fermentations

Minimal media microfermentations were performed as previously reported, ^24,29,30^ except that xylose was substituted for glucose (1 gram xylose for 1 gram glucose) in all media formulations. Guide array stability was confirmed after transformation of pCASCADE plasmids by PCR prior to evaluation according to Li et al.^24^ Fed batch fermentations were performed as previously reported, again with xylose instead of glucose.^30^ Xylose feeding was as modified as follows. The starting batch glucose concentration was 25 g/L. Concentrated sterile filtered xylose feed (500 g/L) was added to the tanks at an initial rate of 10 g/h when cells entered mid-exponential growth. This rate was then increased exponentially, doubling every 1.083 hours (65 min) until 40 g total glucose had been added, after which the feed was maintained at 1.75g/hr. The feed was reduced to 0.875 g/hr due to xylose accumulation at 85 hrs post inoculation, and stopped at 120hrs post inoculation.

### (XyrA) Xylose reductase Purification and Activity assays

*E. coli* BL21(DE3) (New England Biolanbs, Ipswich, MA) with plasmid pHCKan-xyrA (bearing a 6x his tag) was cultured overnight in Luria Broth (Lenox formulation). The overnight culture was used to inoculate SM10++ media (with xylose as a carbon source instead of glucose) with appropriate antibiotics. Cells were cultured at 37°C for 16 hours, then cells were centrifuged and the pellet was washed with SM10 No phosphate media. Next, the washed pellet was resuspended and cultured in SM10 No Phosphate media again with the appropriate antibiotics. After the expression, the post production cells were lysed by a freeze-thaw cycle. XyrA protein was purified using Ni-NTA Resin (G-Biosciences, Cat # 786-939) according to manufacturer’s instructions. Kinetics assays for XyrA were performed in a reaction buffer composed of 50 mM sodium phosphate (pH 7.6, 5mM MgCl2) with NADPH as cofactor. ^55^ In these assays, NADPH was held at a constant initial level of 50 *μ*M. Results of the assay were measured through monitoring the absorbance of NADPH at 340nm for 1.5 hours (15s per read) using a SpectraMax Plus 384 microplate reader (Molecular Devices). Reaction velocity is plotted as a function of xylose concentration. Using the Eadie-hofstee equation, we got the parameters: Vmax=22.6 ± 1.01 U, kcat=13.56 ± 3.05 s^−1^ and Km: 35.12 ± 3.05 mM.

### (XylA) Xylose isomerase quantification

Xylose isomerase activities from cell extracts were quantified with a D-xylose reductase coupled enzyme assay, similar to methods previously described, and following a decrease in absorbance of NADPH at 340nm.^56,57^ Cultures were grown in shake flasks in SM10++ meda and harvested in mid exponential phase, washed and resuspended in SM 10 No phosphate media. After 16 hours of phosphate depletion, cells were pelleted by 10 minutes of centrifugation (4122 RCF, 4 degrees C) and lysed with BugBuster protein extraction reagent (Millipore Sigma, Catalog #70584) according to the manufacturer’s protocol. Cell debris was removed by two rounds of centrifugation, 20 minutes (4122 RCF, 4 degrees C) followed by a 20 minute hard spin (14000 RCF, 4 degrees C). The lysate was filtered with Amicon 30MWCO filters (Millipore Sigma, Catalog #UFC8030) according to the manufacturer’s protocol and washed three times to exchange the buffer with the reaction buffer (45mM sodium phosphate, 10mM MgCl_2_, pH 7.6) and remove metabolites. Samples were assayed in triplicate in a 96 well plate with 100uL of the filtered cell extract per well containing 31.25mM xylulose, 0.5mM NADPH, and 1ug/mL of purified D-xylose reductase (see above). The absorbance at 340nm was measured every 15seconds for 1.5hours and the slope of the linear region was used to quantify XylA activity. Total protein concentration of each sample was determined with a standard Bradford assay. Kinetic parameters were as follows: kcat: 13.56 ± 3.05 s^−1^, Km: 35.12 ± 3.05 mM.

### (UdhA) Soluble transhydrogenase quantification

The activity of the soluble transhydrogenase was quantified by method previously reported. ^28,58^ The process of UdhA expression and cell lysis was carried out using the same method as the XyrA expression mentioned above. The lysates were centrifuged for 15 minutes (4200 RPM, 4°C) to remove large debris. A second hard spin was performed for 30 minutes (14000 RPM, 4°C) to remove remaining debris and further separate the membrane fraction from the soluble transhydrogenase. Lysates were diluted 1:5 with the assay reaction buffer (50mM Tris-HCl, 2mM MgCl, pH 7.6) and transferred to an Amicon Ultra centrifugal filter (10kDa MWCO). The samples were centrifuged for 30 minutes (4200 RPM, 4°C) and this step was repeated 3 times to remove metabolites and exchange the lysis buffer for the assay buffer. After filtration the protein concentrations of the samples were quantified with a standard Bradford assay.

Then soluble transhydrogenase activity was assayed at room temperature. Assays were performed in black 96 well plates by mixing equal volumes of lysate and reaction buffer for a final volume of 100uL per well and a final concentration of 0.5mM NADPH and 1mM 3-acetylpyridine adenine dinucleotide (APAD^+^). Changes in absorbance at 400nm and 310nm due to the reduction of APAD^+^ and the oxidation of NADPH, respectively, were monitored simultaneously by Spectramax Plus 384 microplate reader at 30 second intervals for 30 minutes. A standard curve was used to calculate the molar absorptivity of NADPH (3.04*10^3^ M^−1^ cm^−1^). The molar absorptivity was used to convert the measured slope of the linear region to the change in concentration per minute. The specific activity (Units per mg of total protein) was determined by dividing the change in concentration per minute by the protein concentration.

### FabI Quantification

Quantification of FabI via a C-terminal GFP tags was performed using a GFP quantification kit from AbCam (Cambridge, UK, Cat # ab171581) according to manufacturer’s instructions.

### Xylose and Xylitol Quantification

In micro-fermentations, xylose and xylitol were quantified by commercial bioassays from Megazyme (Wicklow, Ireland, Cat # K-XYLOSE and K-SORB), according to the manufacturer’s instructions. An HPLC method coupled with a refractive index detector was used to quantify both xylose as well as xylitol from instrumented fermentations. Briefly, a Rezex ROA-Organic Acid H^+^ (8%) Analysis HPLC Column (Cat #: #00H-0138-K0, Phenomenex, Inc., Torrance, CA, 300 x 7.8 mme;) was employed for the separation of xylose and xylitol. 5 mM Sulfuric acid as the isocratic mobile phase at a flow rate of 0.5mL/min, ar 55 °C,. A Waters Acquity H-Class UPLC integrated with a Waters 2414 Refractive Index (RI) detector (Waters Corp., Milford, MA. USA) was used for detection. The injection volume of samples and standards was 10 μL. Samples were diluted 20 fold in water in order to be in the linear range (0.01 to 20 g/L). MassLynx v4.1 software was used for all the peak integration and analyses.

### NADPH Pool Quantification

NADPH pools were measured t using an NADPH Assay Kit (AbCam, Cambridge, UK, Cat # ab186031) according to manufacturer’s instructions. Cultures and phosphate depletion were performed as described above for XyrA expression (except there was no xyrA plasmid in the cell). Cells were lysed using the lysis buffer in the assay kit.

### Metabolic Modeling

*In silico* analyses were performed implementing Constraint-based (COBRA) models for *E. coli*, developed employing the COBRApy Python package ^42^ with a previously reported reconstruction ^43^ as a starting point. This curated *E. coli* K-12 MG1655 reconstruction includes 2,719 metabolic reactions and 1,192 unique metabolites. This model was adapted as follows. First, missing reactions and metabolites for xylitol production and export were added as shown in Supplemental Table S5. All reactions, metabolites stoichiometry and identificators were extracted from the BiGG Models database. ^59^ The resulting model was validated for mass balances and metabolite compartment formulas with COBRApy validation methods. ^60^ Once properly balanced, a growth model was created and analyzed.^61^ Specific evaluated conditions and biomass fluxes are shown in Supplemental Table S6.

Next, experimental data obtained from the xylitol micro-fermentations was used to constrain the model. Specific constraints included: i) setting the ratio for pyruvate consumption through Pyruvate Dehydrogenase (PDH) and Pyruvate-flavodoxin Oxidoreductase (*ybdk*), with 10% and 90% of total flux respectively and ii) setting Ferredoxin/flavodoxin reductase to a reversible reaction and iii) using xylose as a sole carbon source with an input flux of 10 mmol/gCDW*hr under minimal media conditions. Finally a set of specific xylitol production strains were constructed and evaluated *in silico* using Flux Balance Analysis (FBA) to obtain xylitol yields, analyze cofactor and/or metabolites of interest as well as production and consumption fluxes. Specific cases that were analyzed included reduction or increased activity of: Zwf, FabI, GltA, XylA, PntAB and UdhA as shown Supplemental Table S7. For each case/condition the following data was obtained: Xylitol yield, NADPH producing and consuming reaction fluxes and escher maps of central metabolism for flux distribution visualization. Finally, major changes in fluxes between the most relevant strains were analyzed.

## Acknowledgements

We would like to acknowledge the following support: DARPA# HR0011-14-C-0075, ONR YIP #N00014-16-1-2558, and DOE EERE grant #EE0007563. Jennifer Hennigan was supported in part by the NIH Biotechnology Training Grant (T32GM008555).

## Author contributions

S. Li constructed plasmids and strains, performed enzyme assays and microfermentations and fermentation studies and related analyses. E.A. Moreb performed microfermentations. Z. Ye constructed plasmids and strains. J.N. Hennigan performed enzyme assays. T. Yang analyzed results from microfermentations and enzyme assays. M.D. Lynch designed and analyzed experiments, constructed strains and performed analytical analyses. All authors wrote revised and edited the manuscript.

## Figures & Captions

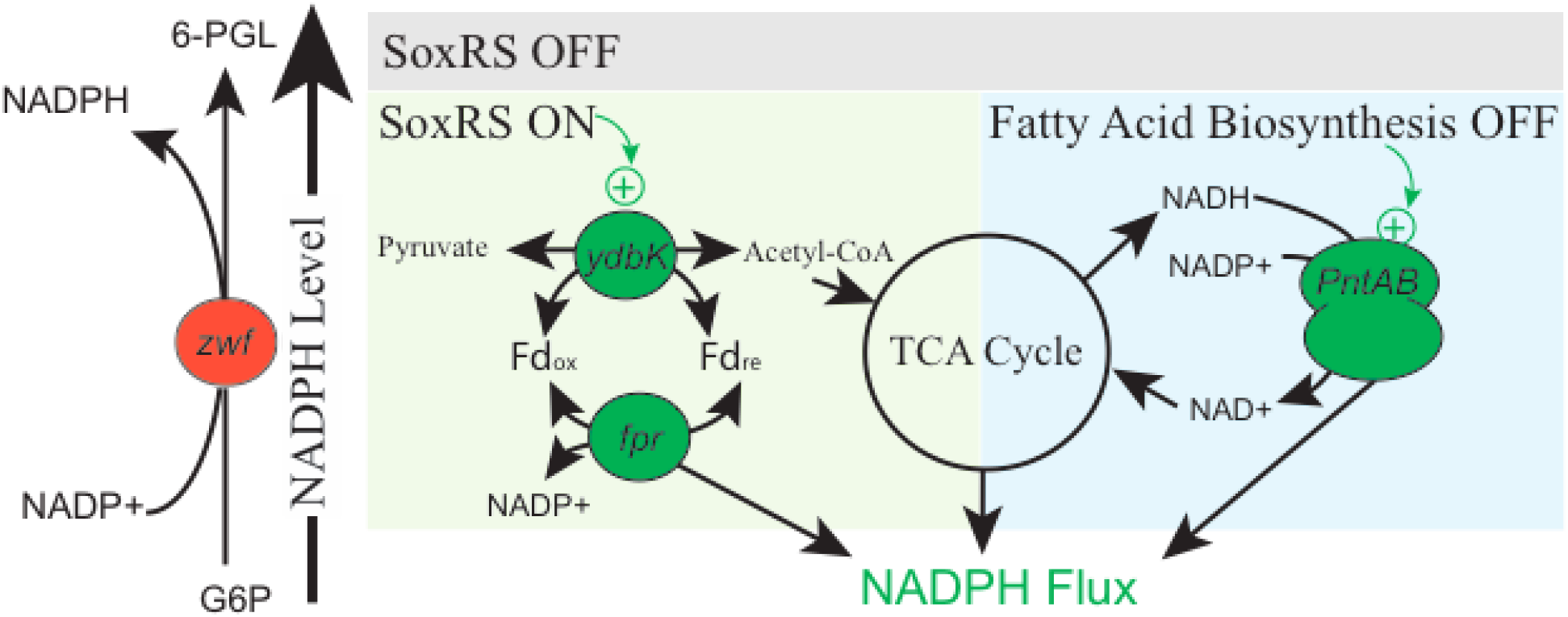
Graphical Abstract.

