## Supplementary Materials for "Dynamic control over feedback regulatory mechanisms improves NADPH fluxes and xylitol biosynthesis in engineered *E. coli*"

**A comparison of stoichiometric and regulatory strategies to improve NADPH fluxes and xylitol biosynthesis, using two-stage dynamic metabolic control in engineered *E. coli***

Shuai Li[1], Eirik A. Moreb [2], Zhixia Ye[2], Jennifer N. Hennigan[1], Daniel Baez Castellanos[2], Tian Yang[2], and Michael D. Lynch [2,3]

### Supplemental Materials

**Table S1.** Plasmids and strains used in this study

| Plasmid | Insert | promoter | Ori | Res | Addgene | Source |
|---|---|---|---|---|---|---|
| pSMART-HC-Kan | None | None | colE1 | Kan | NA | Lucigen |
| pCDF-ev | None | None | cloDF13 | Sm | 89596 | [1] |
| pHCKan-xyrA | xyrA | yibDp[2] | colE1 | Kan | 158610 | This study |
| pCDF-pntAB | pntAB | ugpBp[2] | cloDF13 | Sm | 158609 | This study |
| pCASCADE-ev | none | ugpBp[2] | p15 | Cm | 65821 | [3] |
| pCASCADE-g2 | gltAp2 gRNA | ugpBp[2] | p15 | Cm | 65817 | [3] |
| pCASCADE-f | fabIp gRNA | ugpBp[2] | p15 | Cm | 66635 | This study |
| pCASCADE-z | zwfp gRNA | ugpBp[2] | p15 | Cm | 65825 | [3] |
| pCASCADE-u | udhAp gRNA | ugpBp[2] | p15 | Cm | 65818 | This study |
| pCASCADE-x | xylAp gRNA | ugpBp[2] | p15 | Cm | 158611 | This study |
| pCASCADE-g2z | gltAp2 , zwfp gRNA array | ugpBp[2] | p15 | Cm | 71338 | [3] |
| pCASCADE-g2u | gltAp2 udhAp gRNA array | ugpBp[2] | p15 | Cm | 65819 | This study |
| pCASCADE-g2x | gltAp2, xylAp gRNA array | ugpBp[2] | p15 | Cm | 158613 | This study |

| | | | | | | |
|---|---|---|---|---|---|---|
| pCASCADE-zx | zwfp xylAp gRNA array | ugpBp[2] | p15 | Cm | 158614 | This study |
| pCASCADE-ux | udhAp , xylAp gRNA array | ugpBp[2] | p15 | Cm | 158612 | This study |
| pCASCADE-fg2 | fabIp, gltAp2 gRNA array | ugpBp[2] | p15 | Cm | 71341 | This study |
| pCASCADE-fz | fabIp, zwf gRNA array | ugpBp[2] | p15 | Cm | 71335 | This study |

**Strains used in this study**

| **Strain** | **Genotype** | **Source** |
|---|---|---|
| DLF_Z0025 | F-, λ-, Δ(araD-araB)567, lacZ4787(del)(::rrnB-3) , rph-1, Δ(rhaD-rhaB)568, hsdR514, ΔackA-pta, ΔpoxB, ΔpflB, ΔldhA, ΔadhE, ΔiclR, ΔarcA, ΔsspB, Δcas3::tm-ugpb-sspB-pro-casA. | [3] |
| DLF_Z0043 | DLF_Z0025, gltA-DAS+4-zeoR | [3] |
| DLF_Z1002 | DLF_Z0025, zwf-DAS+4-bsdR | [3] |
| DLF_Z0044 | DLF_Z0025, gltA-DAS+4-zeoR, zwf-DAS+4-bsdR | [3] |
| DLF_Z0028 | DLF_Z0025, fabI-DAS+4-gentR | This study |
| DLF_Z0028G | DLF_Z0025, fabI-sfGFP-gentR | This study |
| DLF_Z0028GD | DLF_Z0025, fabI-sfGFP-DAS+4-gentR | This study |
| DLF_Z0763 | DLF_Z0025, udhA-DAS+4-bsdR | This study |
| DLF_Z0039 | fDLF_Z0025, fabI-DAS+4-gentR, gltA-DAS+4-zeoR | This study |
| DLF_Z0040 | DLF_Z0025, fabI-DAS+4-gentR, zwf-DAS+4-bsdR | This study |
| DLFZ_0045 | DLF_Z0025, fabI-DAS+4-gentR, udhA-DAS+4-bsdR | This study |
| DLFZ_0046 | DLF_Z0025, fabI-DAS+4-gentR, gltA-DAS+4-zeoR, zwf-DAS+4-bsdR | This study |
| DLFZ_0047 | DLF_Z0025, fabI-DAS+4-gentR, gltA-DAS+4-zeoR, udhA-DAS+4-bsdR | This study |
| SL_0001 | DLF_Z0025, xylA-DAS+4-ampR | This study |
| SL_0002 | DLF_Z0025, fabI-DAS+4-gentR, xylA -DAS+4-ampR | This study |

| | | |
|---|---|---|
| SL_0003 | DLF_Z0025, fabI-DAS+4-gentR, gltA-DAS+4-zeoR, xylA -DAS+4-ampR | This study |
| SL_0004 | DLF_Z0025, fabI-DAS+4-gentR, zwf-DAS+4-bsdR, xylA -DAS+4-ampR | This study |
| SL_0005 | DLF_Z0025, fabI-DAS+4-gentR, udhA-DAS+4-bsdR, xylA -DAS+4-ampR | This study |
| SL_0006 | DLF_Z0025, fabI-DAS+4-gentR, gltA-DAS+4-zeoR,udhA-DAS+4-bsdR, xylA -DAS+4-ampR | This study |
| SL_0007 | DLF_Z0025, fabI-DAS+4-gentR, gltA-DAS+4-zeoR, zwf-DAS+4-bsdR, xylA -DAS+4-ampR | This study |
| SL_0008 | DLF_Z0025, gltA-DAS+4-zeoR, udhA-DAS+4-bsdR | This study |
| SL_0009 | DLF_Z0025, zwf-DAS+4,-bsdR xylA -DAS+4-ampR | This study |
| SL_0010 | DLF_Z0025, fabI-DAS+4-gentR, zwf-DAS+4-bsdR, ΔydbK | This study |
| SL_0011 | DLF_Z0025, fabI-DAS+4-gentR, zwf-DAS+4-bsdR, Δfpr | This study |
| Ori- origin of replication, Res - resistance marker, Sm - spectinomycin, Cm- chloramphenicol, Kan - kanamycin, Amp - ampicillin | | |

**Table S2.** Synthetic DNA utilized for strain construction.

| xylA-DAS4-ampR |
|---|
| GATACGATGGCACTGGCGCTGAAAATTGCAGCGCGCATGATTGAAGATGGCGAGCTGGATAAACGC ATCGCGCAGCGTTATTCCGGCTGGAATAGCGAATTGGGCCAGCAAATCCTGAAAGGCCAAATGTCA CTGGCAGATTTAGCCAAATATGCTCAGGAACATCATTTGTCTCCGGTGCATCAGAGTGGTCGCCAGG AACAACTGGAAAATCTGGTAAACCATTATCTGTTCGACAAAGCGGCCAACGATGAAAACTATTCTG AAAACTATGCGGATGCGTCTTAATGATAAGGACCGTGTTGACAATTAATCATCGGCATAGTATATCG GCATAGTATAATACGACAAGGTGAGGAACTAAACCATGAGTATTCAACATTTCCGTGTCGCCCTTAT TCCCTTTTTTGCGGCATTTTGCCTTCCTGTTTTTGCTCACCCAGAAACGCTGGTGAAAGTAAAAGATG CTGAAGATCAGTTGGGTGCACGAGTGGGTTACATCGAACTGGATCTCAACAGCGGTAAGATCCTTG AGAGTTTACGCCCCGAAGAACGTTTTCCAATGATGAGCACTTTTAAAGTTCTGCTATGTGGCGCGGT ATTATCCCGTATTGACGCCGGGCAAGAGCAACTCGGTCGCCGCATACACTATTCTCAGAATGACTTG GTTGAGTACTCACCAGTCACAGAAAAGCATCTCACGGATGGCATGACAGTAAGAGAATTATGCAGT GCTGCCATAACCATGAGTGATAACACTGCGGCCAACTTACTTCTGGCAACGATCGGAGGACCGAAG GAGCTAACCGCTTTTTTGCACAACATGGGGGATCATGTAACTCGCCTTGATCGTTGGGAACCGGAGC TGAATGAAGCCATACCAAACGACGAGCGTGACACCACGATGCCTGTAGCAATGGCAACAACGTTGC GCAAACTATTAACTGGCGAACTACTTACTCTAGCTTCCCGGCAACAATTAATAGACTGGATGGAGGC GGATAAAGTTGCAGGATCACTTCTGCGCTCGGCCCTCCCGGCTGGCTGGTTTATTGCTGATAAATCT |

| |
|---|
| GGAGCCGGTGAGCGTGGGTCTCGCGGTATCATTGCAGCACTGGGGCCAGATGGTAAGCCCTCCCGC<br>ATCGTAGTTATCTACACGACGGGGAGTCAGGCAACTATGGATGAACGAAATAGACAGATCGCTGAG<br>ATAGGTGCCTCACTGATTAAGCATTGGTAGTAAGTAGGGATAACAGGGTAATCGGCTAACTGTGCA<br>GTCCGTTGGCCCGGTTATCGGTAGCGATACCGGGCATTTTTTTAAGGAACGATCGATATGTATATCG<br>GGATAGATCTTGGCACCTCGGGCGTAAAAGTTATTTTGCTCAACGAGCAGGGTGAGGTGGTTGCTGC<br>GCAAACGGAAAAGCTGACCGTTTCGCGCCCGCATCCACTCTGGTCGGAACAAGACCCGGAACAGTG<br>GTGGCAGGCAACTGATCGCGCAA |
| fabI-DAS+4-gentR |
| CTATTGAAGATGTGGGTAACTCTGCGGCATTCCTGTGCTCCGATCTCTCTGCCGGTATCTCCGGTGA<br>AGTGGTCCACGTTGACGGCGGTTTCAGCATTGCTGCAATGAACGAACTCGAACTGAAAGCGGCCAA<br>CGATGAAAACTATTCTGAAAACTATGCGGATGCGTCTTAATAGGAAGTTCCTATTCTCTAGAAAGTA<br>TAGGAACTTCCGAATCCATGTGGGAGTTTATTCTTGACACAGATATTTATGATATAATAACTGAGTA<br>AGCTTAACATAAGGAGGAAAAACATATGTTACGCAGCAGCAACGATGTTACGCAGCAGGGCAGTCG<br>CCCTAAAACAAAGTTAGGTGGCTCAAGTATGGGCATCATTCGCACATGTAGGCTCGGCCCTGACCA<br>AGTCAAATCCATGCGGGCTGCTCTTGATCTTTTCGGTCGTGAGTTCGGAGACGTAGCCACCTACTCC<br>CAACATCAGCCGGACTCCGATTACCTCGGGAACTTGCTCCGTAGTAAGACATTCATCGCGCTTGCTG<br>CCTTCGACCAAGAAGCGGTTGTTGGCGCTCTCGCGGCTTACGTTCTGCCCAAGTTTGAGCAGCCGCG<br>TAGTGAGATCTATATCTATGATCTCGCAGTCTCCGGCGAGCACCGGAGGCAGGGCATTGCCACCGC<br>GCTCATCAATCTCCTCAAGCATGAGGCCAACGCGCTTGGTGCTTATGTGATCTACGTGCAAGCAGAT<br>TACGGTGACGATCCCGCAGTGGCTCTCTATACAAAGTTGGGCATACGGGAAGAAGTGATGCACTTT<br>GATATCGACCCAAGTACCGCCACCTAAGAAGTTCCTATTCTCTAGAAAGTATAGGAACTTCCGTTCT<br>GTTGGTAAAGATGGGCGGCGTTCTGCCGCCCGTTATCTCTGTTATACCTTTCTGATATTTGTTATCGC<br>CGATCCGTCTTTCTCCCCTTCCCGCCTTGCGTCAGG |
| fabI-sfGFP-gentR |
| AAAGACTTCCGCAAAATGCTGGCTCATTGCGAAGCCGTTACCCCGATTCGCCGTACCGTTACTATTG<br>AAGATGTGGGTAACTCTGCGGCATTCCTGTGCTCCGATCTCTCTGCCGGTATCTCCGGTGAAGTGGT<br>CCACGTTGACGGCGGTTTCAGCATTGCTGCAATGAACGAACTCGAACTGAAAGGGGGTTCAGGCGG<br>GTCGGGTGGCgtgagcaagggcgaggagctgttcaccggggtggtgcccatcctggtcgagctggacggcgacgtaaacggccacaagttcagcgtgc<br>gcggcgagggcgagggcgatgccaccaacggcaagctgaccctgaagttcatctgcaccaccggcaagctgcccgtgccctggcccaccctcgtgaccaccctg<br>acctacggcgtgcagtgcttcagccgctaccccgaccacatgaagcgccacgacttcttcaagtccgccatgcccgaaggctacgtccaggagcgcaccatcagct<br>tcaaggacgacggcacctacaagacccgcgccgaggtgaagttcgagggcgacaccctggtgaaccgcatcgagctgaagggcatcgacttcaaggaggacgg<br>caacatcctggggcacaagctggagtacaacttcaacagccacaacgtctatatcaccgccgacaagcagaagaacggcatcaaggccaacttcaagatccgcca<br>caacgtggaggacggcagcgtgcagctcgccgaccactaccagcagaacacccccatcggcgacggccccgtgctgctgcccgacaaccactacctgagcacc<br>cagtccgtgctgagcaaagaccccaacgagaagcgcgatcacatggtcctgctggagttcgtgaccgccgccgggatcactcacggcatggacgagctgtacaag<br>TAATGACGAATCCATGTGGGAGTTTATTCTTGACACAGATATTTATGATATAATAACTGAGTAAGCT<br>TAACATAAGGAGGAAAAACATATGTTGCGTAGCTCTAACGATGTGACGCAACAAGGTTCGCGTCCA<br>AAGACAAAATTGGGAGGCAGTAGCATGGGGATCATTCGCACTTGTCGCCTGGGGCCAGACCAGGTG<br>AAGTCAATGCGTGCGGCTCTGGACTTATTCGGGCGCGAATTTGGAGATGTAGCCACTTACTCACAGC<br>ACCAACCGGACAGTGATTACTTGGGGAATTTACTTCGCAGTAAAACTTTTATCGCTTTGGCCGCTTT<br>CGACCAGGAGGCTGTAGTAGGTGCGTTGGCAGCCTATGTTCTTCCTAAATTCGAGCAACCGCGTAGC<br>GAAATTTACATCTATGATCTTGCAGTCTCCGGCGAACATCGCCGTCAGGGGATCGCCACAGCTTTAA<br>TCAACCTTTTGAAGCATGAGGCTAATGCACTTGGAGCGTACGTGATTTATGTGCAGGCTGACTACGG<br>TGATGATCCTGCAGTCGCTCTGTACACCAAACTGGGTATCCGCGAGGAGGTCATGCACTTTGATATT |

| |
|---|
| GACCCGTCGACGGCTACCTAAGTTCTGTTGGTAAAGATGGGCGGCGTTCTGCCGCCCGTTATCTCTG<br>TTATACCTTTCTGATATTTGTTATCGCCGATCCGTCTTTCTCCCCTTCCCGCCTTGCGTCAGG |
| fabI-sfGFP-DAS+4- gentR |
| AAAGACTTCCGCAAAATGCTGGCTCATTGCGAAGCCGTTACCCCGATTCGCCGTACCGTTACTATTG<br>AAGATGTGGGTAACTCTGCGGCATTCCTGTGCTCCGATCTCTCTGCCGGTATCTCCGGTGAAGTGGT<br>CCACGTTGACGGCGGTTTCAGCATTGCTGCAATGAACGAACTCGAACTGAAAGGGGGTTCAGGCGG<br>GTCGGGTGGCgtgagcaagggcgaggagctgttcaccggggtggtgcccatcctggtcgagctggacggcgacgtaaacggccacaagttcagcgtgc<br>gcggcgagggcgagggcgatgccaccaacggcaagctgaccctgaagttcatctgcaccaccggcaagctgcccgtgccctggcccaccctcgtgaccaccctg<br>acctacggcgtgcagtgcttcagccgctaccccgaccacatgaagcgccacgacttcttcaagtccgccatgcccgaaggctacgtccaggagcgcaccatcagct<br>tcaaggacgacggcacctacaagacccgcgccgaggtgaagttcgagggcgacaccctggtgaaccgcatcgagctgaagggcatcgacttcaaggaggacgg<br>caacatcctggggcacaagctggagtacaacttcaacagccacaacgtctatatcaccgccgacaagcagaagaacggcatcaaggccaacttcaagatccgcca<br>caacgtggaggacggcagcgtgcagctcgccgaccactaccagcagaacacccccatcggcgacggccccgtgctgctgcccgacaaccactacctgagcacc<br>cagtccgtgctgagcaaagaccccaacgagaagcgcgatcacatggtcctgctggagttcgtgaccgccgccgggatcactcacggcatggacgagctgtacaag<br>GGTGGGGGTGGGAGCGGCGGCGGTGGCTCCGCGGCCAACGATGAAAACTATTCTGAAAACTATGCG<br>GATGCGTCTTAATGACGAATCCATGTGGGAGTTTATTCTTGACACAGATATTTATGATATAATAACT<br>GAGTAAGCTTAACATAAGGAGGAAAAACATATGTTGCGTAGCTCTAACGATGTGACGCAACAAGGT<br>TCGCGTCCAAAGACAAAATTGGGAGGCAGTAGCATGGGGATCATTCGCACTTGTCGCCTGGGGCCA<br>GACCAGGTGAAGTCAATGCGTGCGGCTCTGGACTTATTCGGGCGCGAATTTGGAGATGTAGCCACTT<br>ACTCACAGCACCAACCGGACAGTGATTACTTGGGGAATTTACTTCGCAGTAAAACTTTTATCGCTTT<br>GGCCGCTTTCGACCAGGAGGCTGTAGTAGGTGCGTTGGCAGCCTATGTTCTTCCTAAATTCGAGCAA<br>CCGCGTAGCGAAATTTACATCTATGATCTTGCAGTCTCCGGCGAACATCGCCGTCAGGGGATCGCCA<br>CAGCTTTAATCAACCTTTTGAAGCATGAGGCTAATGCACTTGGAGCGTACGTGATTTATGTGCAGGC<br>TGACTACGGTGATGATCCTGCAGTCGCTCTGTACACCAAACTGGGTATCCGCGAGGAGGTCATGCAC<br>TTTGATATTGACCCGTCGACGGCTACCTAAGTTCTGTTGGTAAAGATGGGCGGCGTTCTGCCGCCCG<br>TTATCTCTGTTATACCTTTCTGATATTTGTTATCGCCGATCCGTCTTTCTCCCCTTCCCGCCTTGCGTC<br>AGG |
| udhA-DAS+4-bsdR |
| TCTGGGTATTCACTGCTTTGGCGAGCGCGCTGCCGAAATTATTCATATCGGTCAGGCGATTATGGAA<br>CAGAAAGGTGGCGGCAACACTATTGAGTACTTCGTCAACACCACCTTTAACTACCCGACGATGGCG<br>GAAGCCTATCGGGTAGCTGCGTTAAACGGTTTAAACCGCCTGTTTGCGGCCAACGATGAAAACTATT<br>CTGAAAACTATGCGGATGCGTCTTAATAGTTGACAATTAATCATCGGCATAGTATATCGGCATAGTA<br>TAATACGACTCACTATAGGAGGGCCATCATGAAGACCTTCAACATCTCTCAGCAGGATCTGGAGCT<br>GGTGGAGGTCGCCACTGAGAAGATCACCATGCTCTATGAGGACAACAAGCACCATGTCGGGGCGGC<br>CATCAGGACCAAGACTGGGGAGATCATCTCTGCTGTCCACATTGAGGCCTACATTGGCAGGGTCACT<br>GTCTGTGCTGAAGCCATTGCCATTGGGTCTGCTGTGAGCAACGGGCAGAAGGACTTTGACACCATTG<br>TGGCTGTCAGGCACCCCTACTCTGATGAGGTGGACAGATCCATCAGGGTGGTCAGCCCCTGTGGCAT<br>GTGCAGAGAGCTCATCTCTGACTATGCTCCTGACTGCTTTGTGCTCATTGAGATGAATGGCAAGCTG<br>GTCAAAACCACCATTGAGGAACTCATCCCCCTCAAGTACACCAGGAACTAAAGTAAAACTTTATCG<br>AAATGGCCATCCATTCTTGCGCGGATGGCCTCTGCCAGCTGCTCATAGCGGCTGCGCAGCGGTGAGC<br>CAGGACGATAAACCAGGCCAATAGTGCGGCGTGGTTCCGGCTTAATGCACGG |

**Table S3:** Oligos used for strain confirmations & sequencing

| Plasmids/primers name | Sequences |
|---|---|
| fabI_int_F | GCAAAATGCTGGCTCATTG |
| gentR_intR | GCGATGAATGTCTTACTACGGA |
| gltA_int_F | TATCATCCTGAAAGCGATGG |
| zeo_intR | ACTGAAGCCCAGACGATC |
| zwf_intF | CTGCTGGAAACCATGCG |
| udhA_intF | CAAAAGAGATTCTGGGTATTCACT |
| bsdR_intR | GAGCATGGTGATCTTCTCAGT |
| xylA_intF | AGATGGCGAGCTGGATA |
| ampR_intR | AGTACTCAACCAAGTCATTCTG |

**Table S4:** Oligos used for pCASCADE plasmid construction

| Plasmids/primers name | Sequences |
|---|---|
| gltA2 | TATTGACCAATTCATTCGGGACAGTTATTAGTTCGAGTTCCCCGCGCCAGCGGGGATAAACCG |
| gltA2-FOR | CCGGATGAGCATTCATCAGGCGGGCAAG |
| gltA2-REV | CGGTTTATCCCCGCTGGCGCGGGGAACTCGAACTAATAACTGTC |
| udhA | TTACCATTCTGTTGCTTTTATGTATAAGAATCGAGTTCCCCGCGCCAGCGGGGATAAACCG |
| udhA-FOR | CCGGATGAGCATTCATCAGGCGGGCAAG |
| udhA-REV | CGGTTTATCCCCGCTGGCGCGGGGAACTCGATTCTTATACATAAAAGC |
| xylA | GGAGTGCCCAATATTACGACATCATCCATCTCGAGTTCCCCGCGCCAGCGGGGATAAACCG |

| | |
|---|---|
| xylA-FOR | CCAGCGGGGATAAACCGGGAGTGCCCAATATTAC |
| xylA-REV | CTTGCCCGCCTGATGAATGCTCATCCGG |

**Table S5.** Xylitol Specific reactions added to the metabolic model

| Name | Reaction | Identifier |
|---|---|---|
| Xylose reductase | h_c + nadph_c + xyl__D_c ⇌ nadp_c + xylt_c | XYLR |
| Xylitol exchange | xylt_e ⇌ | EX_xylt_e |
| Xylitol transport via passive diffusion | xylt_e ⇌ xylt_c | XYLTt |
| Flavodoxin reductase (NADPH) | 2.0 flxso_c + nadph_c ⇌ 2.0 flxr_c + h_c + nadp_c | FLDR2 |

**Table S6.** Metabolic model validation with different carbon sources and oxygen levels. (All flux values are expressed in nmol/gDW*hr).

| Only Carbon Source | C.S Flux | Oxygen Flux | Biomass Flux obtained from model | Previously reported Biomass Flux* |
|---|---|---|---|---|
| Glucose | -8.8 | -30 (Aerobic) | 0.771914 | 0.70± 0.01 |
| Glucose | -8.8 | 0 (Anaerobic) | 0.253285 | 0.33 ± 0.02 |
| Xylose | -9.5 | -30 (Aerobic) | 0.645282 | 0.50 ± 0.02 |
| Xylose | -9.5 | 0 (Anaerobic) | 0.115445 | 0.13 ± 0.02 |

*Numbers taken from [4]

**Table S7.** Strains modeled

| Individual Valves | | Combinations of Valves | |
|---|---|---|---|
| Strain | Valves Silencing | Strain | Valves Silencing |
| WT | -- | Z-F | zwf & fabI |
| Z | zwf | Z-G | zwf & gltA |

| | | | |
|---|---|---|---|
| F | fabI | Z-X | zwf & xylA |
| G | gltA | Z-pAB | zwf & pntAB |
| X | xylA | Z-F-G | zwf, fabI & gltA |
| pAB | pntAB | Z-G-U | zwf, gltA & udhA |
| U | udhA | Z-F-pAB | zwf, fabI & pntAB |

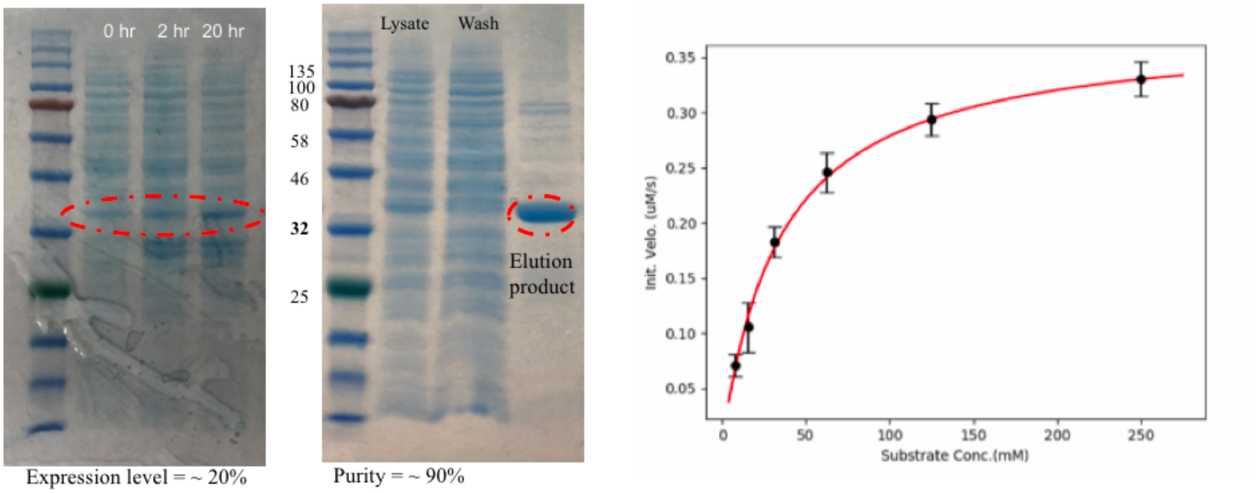


**Figure S1:** XyrA expression and purification from BL21(DE3). Left: A time course of expression post phosphate depletion, whole cell lysates demonstrate expression of XyrA. Densitometry indicates an expression level of ~ 20%. Middle: Purification of XyrA (which contains an N-terminal 6 X histidine tag) via IMAC. Right Kinetic analysis of purified XyrA. Initial velocity ($\mu$M/s) is plotted as a function of substrate (xylose) concentration.

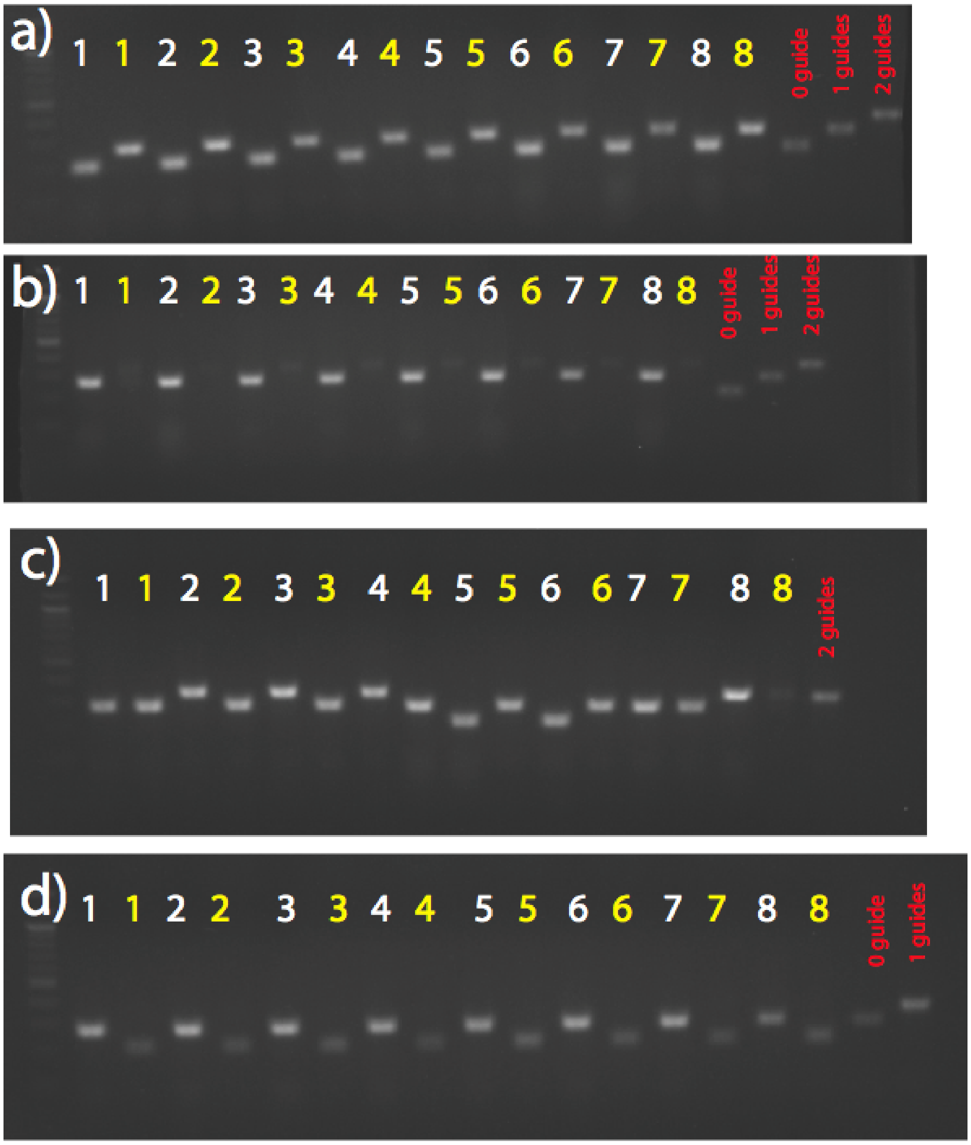


**Figure S2:** Agarose gel electrophoretic analysis of gRNA array stability. Colony PCR was used to amplify and size gRNA arrays from 8 clones after transformation into host strains engineered for dynamic metabolic control. Red indicates PCR products are taken from sequence confirmed gRNA arrays with 0, 1 , or 2 gRNAs respectively. a) Strain DLF_Z0025, white labels: pCASCADE-ev, yellow labels: pCASCADE-g2, b) Strain DLF_Z0025, white labels: pCASCADE-z, yellow labels: pCASCADE-fg2, c) White labels: Strain DLF_Z0044, pCASCADE-fg2, yellow labels: DLF_Z0025, pCASCADE-fg2, d) White labels: Strain DLF_Z0046 , pCASCADE-g2, yellow labels: DLF_Z1002 pCASCADE-fg2.

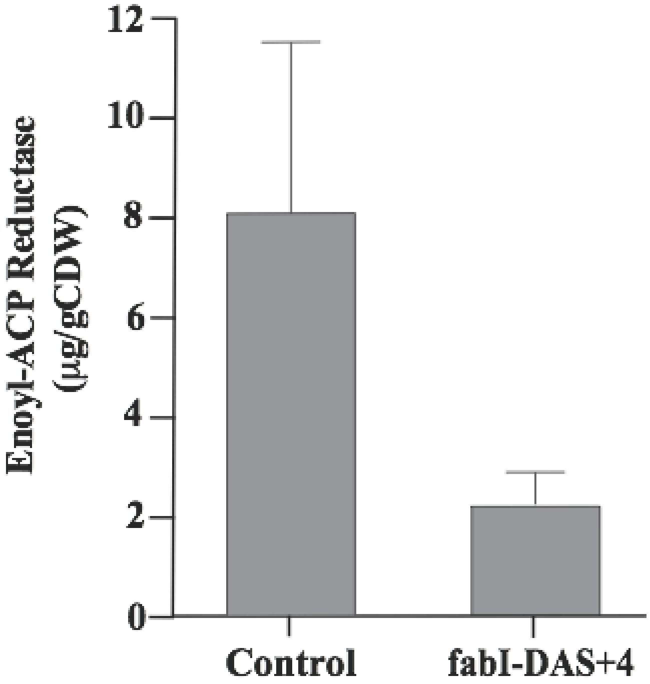


**Figure S3:** Dynamic Control over FabI (enoyl-ACP reductase) levels due to induble proteolysis with a DAS+4 degron tag. The chromosomal fabI gene was tagged with a C-terminal sfGFP. Protein levels were measured by ELISA, 24 hour post induction by phosphate depletion in microfermentations.

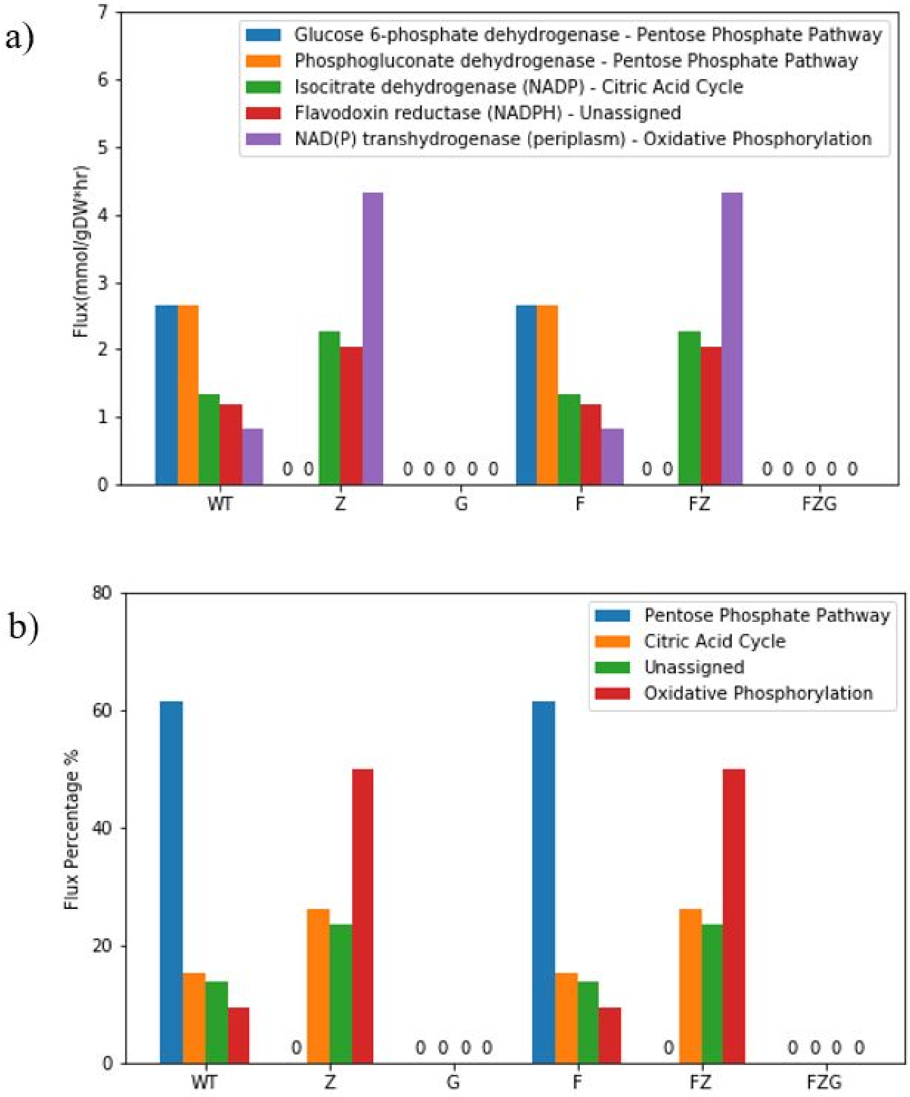


**Figure S4:** Modelled NADPH producing reactions and pathways for Xylitol production in different production strains. a) Specific reactions fluxes during xylitol production. b) Pathway percentage fluxes for xylitol production.
